# Loss of PRMT2 in myeloid cells in normoglycemic mice phenocopies impaired regression of atherosclerosis in diabetic mice

**DOI:** 10.1101/2022.04.15.488480

**Authors:** Beyza Vurusaner, Prashanth Thevkar-Nages, Ravneet Kaur, Chiara Giannarelli, Michael J. Garabedian, Edward A. Fisher

## Abstract

Regression of atherosclerosis is impaired in diabetes. However, the factors mediating this effect remain incomplete. We identified Protein Arginine Methyltransferase 2 (PRMT2) as a protein whose expression in macrophages is reduced in hyperglycemia and diabetes. PRMT2 catalyzes arginine methylation to target proteins to modulate gene expression. Because PRMT2 expression is reduced in cells in hyperglycemia, we wanted to determine whether PRMT2 plays a causal role in the impairment of atherosclerosis regression in diabetes. We therefore examined the consequence of deleting PRMT2 in myeloid cells during regression of atherosclerosis in normal and diabetic mice. Remarkably, we found a significant impairment of atherosclerosis regression under normoglycemic conditions in mice lacking PRMT2 (*Prmt2^-/-^*) in myeloid cells that mimics the decrease in regression of atherosclerosis in WT mice under diabetic conditions. This was associated with increased plaque macrophage retention, as well as increased apoptosis and necrosis. PRMT2-deficient plaque CD68+ cells under normoglycemic conditions showed increased expression of genes involved in cytokine signaling and inflammation compared to WT. Consistently *Prmt2^-/-^* bone marrow derived macrophages (BMDMs) showed an increased response of pro-inflammatory genes to LPS, and a decreased response of inflammation resolving genes to IL-4. This increased response to LPS in *Prmt2^-/-^* BMDMs is via enhanced NF-kappa B activity. Thus, the loss of PRMT2 is causally linked to impaired atherosclerosis regression via a heightened inflammatory response in macrophages. That *PRMT2* expression was lower in myeloid cells in plaques from human subjects with diabetes supports the relevance of our findings to human atherosclerosis.

## INTRODUCTION

People with diabetes have more coronary heart disease (CHD) than people without, even though the statin drugs are equally effective in both groups at lowering the blood levels of LDL cholesterol (LDL-C)^1^. Part of the etiology of CHD stems from cholesterol-engorged myeloid-derived macrophages, termed foam cells, that accumulate in arteries and form plaques. These cells have a diminished capacity to efflux cholesterol, migrate and resolve inflammation^2^, issues that appear to be more pronounced in diabetes^3^. This can ultimately lead to the plaque becoming unstable and rupturing, resulting in heart attacks and stroke. Therefore, an important clinical goal to reduce CHD risk is to promote the regression of atherosclerosis by returning the foam cells to a healthier state by restoration of the aforementioned capacities^4,5^.

In diabetes, this has been particularly challenging, as the molecular mechanisms underlying the attenuation of CHD reduction by LDL-C lowering are incompletely determined. To narrow this gap in understanding, we have identified Protein Arginine Methyltransferase 2 (PRMT2) as a protein whose expression in macrophages was extremely low in high-glucose compared to normal glucose conditions^6^. PRMT2 is an enzyme that catalyzes the transfer of a methyl group from S-adenosylmethionine (SAM) asymmetrically to the arginine residues on histones, such as histone H3R8^7,8^, and other proteins to affect their functions^9–11^. In fibroblasts, PRMT2 has been reported to inhibit NF-kappa B-dependent transcription by increasing the nuclear accumulation of the Ikappa B-alpha^12^ which is a repressor of NF-kappa B. It has also been reported that a reduction in PRMT2 copy number in lung macrophages increased their LPS responsiveness and promoted inflammatory cytokine expression^13^. Thus, among other effects, reducing PRMT2 expression has the potential to promote inflammation.

Our group has also reported that deficiency of PRMT2 was associated with high-glucose inhibition of the LXR-dependent expression of LXR target genes involved in the anti-atherosclerotic macrophage cholesterol efflux and reverse cholesterol transport, including ATP-binding cassette transporter A1 (*Abca1*) and apolipoprotein E (*Apoe*)^6^. LXRs are nuclear receptors that control the expression of genes involved in cholesterol homeostasis, and inflammation and are activated by oxidized cholesterol ligands^14^. We also found in mice the expression of LXRα was induced in plaque macrophages undergoing atherosclerosis regression^15^ and is required for atherosclerosis regression after reversal of hypercholesterolemia^16^.

Because we have been able to show that diabetes impaired atherosclerosis regression in a mouse model^17^ in a manner consistent with the clinical data cited, we decided to adapt this model to the study of PRMT2. Given that PRMT2 has the potential to repress inflammation and increase cholesterol efflux pathways through LXR, we hypothesized that the reduced expression of PMRT2 by high glucose is a key determinant of impaired atherosclerosis regression in diabetes. To test this, we examined whether the myeloid-specific deletion of *Prmt2* would impair regression of atherosclerosis in normoglycemic conditions, thus mimicking the decrease in regression of atherosclerosis in WT mice under diabetic conditions. In fact, we found that the deletion of PRMT2 in myeloid cells in normoglycemic conditions phenocopied the impaired atherosclerosis regression in hyperglycemic mice by reducing the ability of macrophages to migrate out of the plaque. This was associated with increases in macrophage (CD68) gene expression linked to cytokine signaling and inflammation. Studies in macrophages *in vitro* implicate effects on NF-kappa B-mediated inflammation. Thus, expression of PRMT2 is a key factor in promoting atherosclerosis regression, and conversely when reduced contributes to impaired atherosclerosis regression. Approaches to increase PRMT2 expression and enhance LXR activity in diabetes/hyperglycemia could facilitate atherosclerosis regression and reduce CHD.

## METHODS

### Mouse Studies

All experimental procedures were done in accordance with the US Department of Agriculture Animal Welfare Act and the US Public Health Service Policy on Humane Care and Use of Laboratory Animals and were approved by the New York University School of Medicine’s Institutional Animal Care and Use Committee. The studies with mice reported conform to the ARRIVE guidelines. To power our studies, sample size was predefined as n=8 to 10 mice/group. Bone marrow from *WT and Prmt2^-/-^* mice^12^ were transplanted into 8 week old *Ldlr^-/-^* mice. After 4 weeks of recovery, the mice were placed on Western diet (21 % [wt/wt] fat, 0.3% cholesterol [Research Diets]) for 20 weeks to allow development of atherosclerotic plaques. Mice were injected IP with STZ (50mg/kg, Sigma-Aldrich) or citrate buffer for 5 days to induce diabetes mellitus or to serve as a control. Regression was induced by reversal of hyperlipidemia using an apoB-antisense oligonucleotide (ASO) (Ionis Pharmaceuticals, 50 mg/kg; twice a week for 3 weeks) on chow diet (13% fat kcal, 0% cholesterol). Baseline groups were sacrificed at this time. Mice were randomly assigned to either baseline or 1 of 4 regression groups: apoB-ASO + citrate (non-diabetic) or apoB-ASO + STZ (diabetic) harboring bone marrow from either WT or *Prmt2^-/-^* donors.

Regression groups were sacrificed after 3 weeks on chow diet. Mice were anesthetized with xylazine/ketamine, and blood was collected via cardiac puncture for plasma analyses. Mice were perfused with 10% sucrose/saline. Aortic roots were dissected and embedded in optimal cutting temperature compound medium (OCT) and frozen immediately and stored at −80°C until further use.

### Labeling and Tracking of Blood Monocytes

Circulating blood monocytes were labeled *in vivo* by retroorbital intravenous injection of 1 μm Flouresbrite YG microspheres (Polysciences Inc, PA) diluted 1:4 in sterile PBS in mice undergoing regression 3 days prior to harvest. The efficiency of bead labeling was verified 24 hours later by flow cytometry. The number of labeled macrophages (Ly6C^lo^) remaining in the aortic root lesions was determined as described^4^. Ly6C^hi^ monocytes were labeled by intraperitoneal injection of 4 mg/mL of Edu (5-ethynyl-2’-deoxyuridine) from Life technologies (NY), and mice were euthanized after 5 days to assess baseline recruitment or 21 days to assess for retention. Efficiency of Edu labeling was assessed using Click-IT EdU Pacific Blue Flow Cytometry Assay Kit (Life Technologies, NY) after 24 hours, and EdU labeled cells were stained using a Click-IT reaction with Alexa Fluor 647 nm-azide (Click-iT EdU Imaging Kit, Invitrogen, CA).

### Tissue Collection and Immunohistochemistry

The aortic roots of mice were harvested at baseline, or 3 weeks after apoB-ASO. Hearts were sectioned through the aortic root (6 μm) and stained with hematoxylin and eosin for quantification of lesion area. For immunostaining, slides were fixed with 4% formaldehyde, permeabilized with 0.1% triton, and blocked with 5% BSA. The sections were incubated with antibodies against CD68 (cat # MCA1957, Bio-Rad, CA) to detect macrophages Ki67 (cat # Ab1666, Abcam, MA) to detect proliferating cells, and cleaved caspase 3 (cat # 9664, Cell Signaling, MA) to detect apoptosis. Sections were then incubated with appropriate secondary antibodies and stained with DAPI to detect nuclei. To distinguish bona fide target staining from background secondary antibody only was used as a control. For collagen quantification, slides were stained with picrosirius red as previously reported^18^, and collagen was visualized using polarizing light microscopy^19^. The images were taken with a Nikon Eclipse microscope and analyzed using Image J software. Quantification of immunostaining was performed from 6 high power fields per aortic root or arch section/mouse. Representative images were selected to represent the mean value of each condition.

### Cell Culture

BMDMs were isolated from the tibia and femur of 8-12-week-old male wild type C57BL6J and *Prmt2^-/-^* mice. Isolated bone marrow cells were treated with red blood cell lysis buffer (Sigma) and re-suspended in differentiation medium (DMEM and L-glutamine with 1 g/L d-glucose + 3.5 g/L l-glucose (normal glucose; NG), or 4.5 g/L d-glucose (high glucose; HG), supplemented with 20% FBS and 10 ng/μL macrophage colony-stimulating factor (M-CSF) (PeproTech, Inc., Rocky Hill, NJ). Cells were passed through a 70 μm filter to clear debris. Following this, cells were plated in 10cm non-tissue coated plates and allowed to differentiate in either NG or HG containing media for 7 days to obtain non-activated (M0) macrophages. At day 7, the cells were washed in PBS, and re-plated at the desired cell density in a 6-well dish in normal and high glucose media and allowed to attach to the plate. Cells were treated with IL-4 (20ng/ml) or (LPS (100ng/ml) for the indicated times, and RNA was isolated. For some experiments, NF-kappa B inhibitor, caffeic acid phenethyl ester [CAPE (5 μM)] or TPCA1 (5μM) was pretreated for 2 hours before LPS treatment.

### Plasma cholesterol and blood glucose analyses

Total cholesterol was measured using colorimetric assays (Wako Diagnostics, Richmond, VA). For glucose measurements, mice were fasted for 4 h, and glucose in tail blood was measured using a blood glucose monitor (TrueTrack Smart System, Nipro Diagnostics, Inc., Fort Lauderdale, FL).

### Laser capture microdissection and RNA sequencing

CD68+ cells were selected from atherosclerotic plaques by laser capture microdissection (LCM) as previously described^20^. All LCM procedures were performed under RNase-free conditions. Aortic root sections were stained with hematoxylin-eosin. Foam cells were identified under a microscope and verified by positive CD68 staining. For each animal, CD68+ cells were captured from 50–60 frozen sections. RNA was isolated using the PicoPure Kit (Molecular Devices, Inc., Sunnyvale, CA), and quality and quantity were determined using an Agilent 2100 Bioanalyzer (Agilent Technologies, Santa Clara, CA). RNA seq libraries were prepared using the Clontech SMARTer Stranded Total RNA Seq Kit -Pico Input Mammalian following the manufacturer’s protocol. Libraries were purified using AMPure beads, pooled equimolarly, and run on a HiSeq 4000 paired end reads. FASTQ files were obtained, and low-quality bases as well as adapter sequences were trimmed using Cutadapt 1.18. Reads were subsequently mapped to the Mus musculus GRCm38 transcriptome using Kallisto 0.46.2. Raw counts per transcript were summed up at the gene level, and differential expression analysis was performed using edgeR 3.28.0. Genes with a *P* < 0.01 and logFC > 2.0 were determined to be differentially expressed. Pathways analysis was performed using Metascape^21^. RNA sequencing and differential gene expression analysis was performed by the NYU Genome Technology Center.

### Data sharing

The RNA seq datasets generated and/or analyzed during the current study will be available in the NCBI GEO repository. Additional data generated and/or analyzed during the current study will be available from the corresponding authors on reasonable request.

### Differential gene expression analysis

*PRMT2* expression in myeloid cells from human atherosclerotic plaques between two conditions (diabetic and non-diabetic) was performed using the Seurat function FindMarkers. The Wilcoxon rank-sum test (two-sided) was used to identify *PRMT2* expression between two groups and the logfc.threshold was set to 0.25, and min.cells.group = 25 (minimum of 25% of cells in any group being compared). P values were adjusted using Bonferroni correction <0.05.

## RESULTS AND DISCUSSION

### PRMT2 in atherosclerosis regression: study design and metabolic parameters

To determine the role of PRMT2 in myeloid cells in atherosclerosis regression, we performed bone marrow transplants from littermate WT and *Prmt2^-/-^* mice into lethally irradiated *Ldlr^-/-^* mice, as diagrammed in Figure 1A. After bone marrow transplant, mice were allowed to recover for 4 weeks and then fed a western diet (WD) for 20 weeks to permit advanced atherosclerotic plaques to develop. Mice were separated into 6 groups after the dietary period: 2 baseline groups of *Ldlr^-/-^* mice that received either wild type or *Prmt2*-deficient bone marrow (designated as *Ldlr^-/-^:WT* and *Ldlr^-/-^:Prmt2^-/-^*, respectively), and four regression groups: 1. normoglycemic WT (NG *Ldlr^-/-^:WT*); 2. normoglycemic *Prmt2^-/-^* (NG *Ldlr^-/-^:Prmt2^-/-^*); 3. Diabetic WT (D *Ldlr^-/-^:WT*); and, 4) Diabetic *Prmt2^-/-^* (D *Ldlr^-/-^:Prmt2^-/-^*). All regression groups had their apoB-lipoprotein cholesterol levels reduced by injection of an ApoB anti-sense oligonucleotide (ASO) (as done before ^22^) and were also switched to a standard chow diet to maximally promote regression. In addition, after 19 weeks of WD feeding, to induce diabetes *Ldlr^-/-^:WT* and *Ldlr^-/-^:Prmt2^-/-^* mice were given daily intraperitoneal injections of STZ for 5 consecutive days (NG mice were injected with citrate buffer) ^17^.

**Figure 1.**
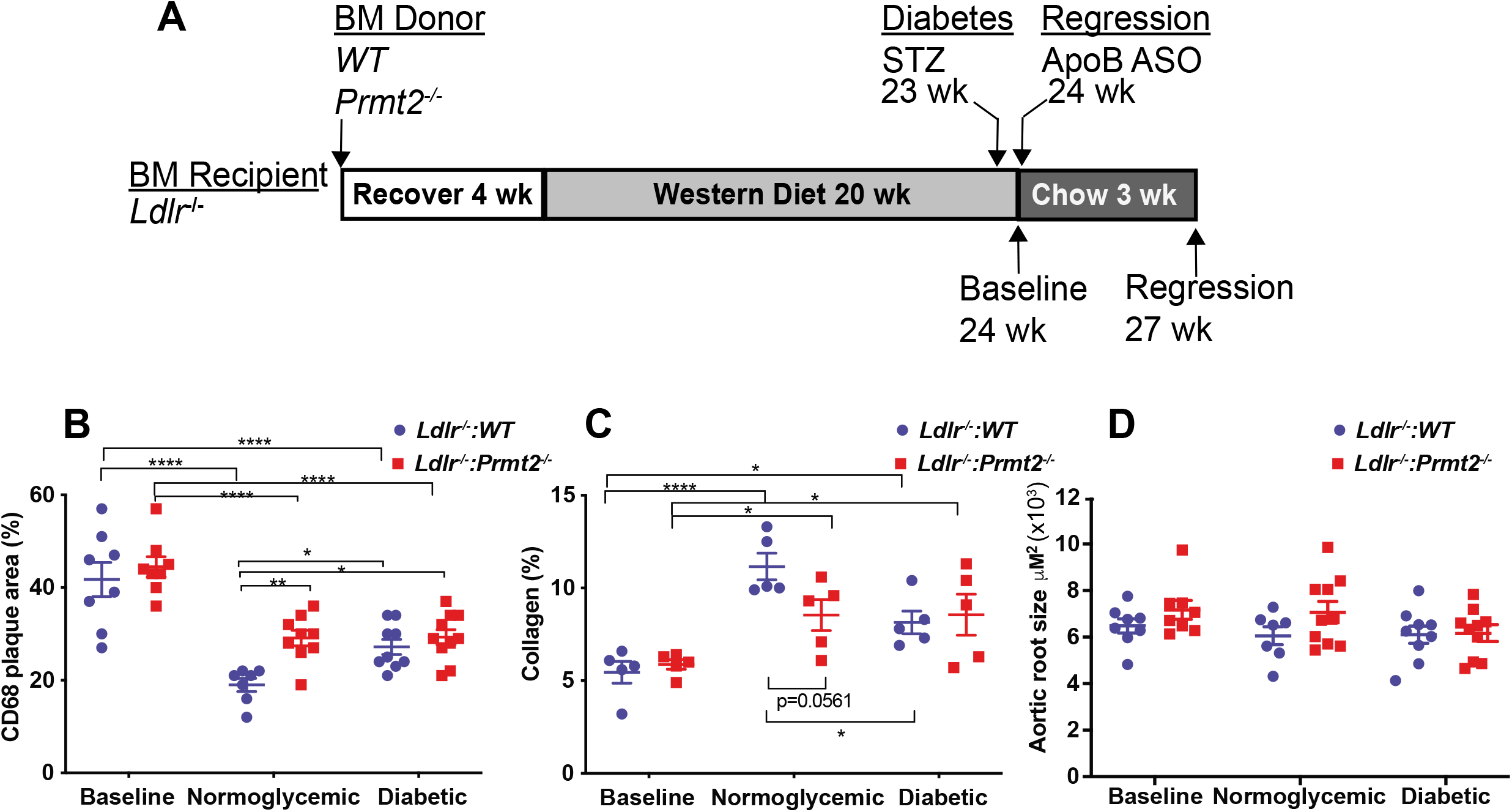
Myeloid PRMT2 deficient mice have increased CD68+ cell and decreased collagen content in nondiabetic conditions after plasma lipid reduction. A) Experimental design: Bone marrow (BM) from WT and *Prmt2^-/-^* donor mice were transplanted into *Ldlr^-/-^* recipient mice. After 4 weeks of recovery mice were placed on a western diet for 20 weeks. A set of animals at after 20 weeks of western diet feeding were used as the baseline group. Two other groups of animals received citrate buffer or STZ at 23 weeks. At 24 weeks, to lower lipids and promote regression, mice received apoB-ASO injections, switched to a chow diet for 3 weeks, and used as the regression groups. B) Aortic roots from baseline and the four regression groups were sectioned and stained for CD68+ cells. Data are shown as the percentage of plaque area. C) Collagen content was determined by picrosirius red staining using both bright field and polarized light microscopy and quantified by Image Pro Plus software. Data are presented as the percentage of lesion area. D) Total plaque areas were quantified by Image Pro Plus software. Each group contains 8-10 animals. Data (means ± SEM) were analyzed using one-way ANOVA followed by Bonferroni’s multiple comparison test. *P* values are shown as \**P* < 0.05 and \*\*\*\**P* < 0.001.

After 19 weeks of WD, the total plasma cholesterol, fasting glucose levels, and body weights (Sup. Fig.1A-C) were similar in the *Ldlr^-/-^* mice reconstituted with WT and *Prmt2^-/-^* bone marrow, as well as the mice randomized into the regression groups (prior to their treatments). ApoB-ASO treatment resulted in significantly lower plasma cholesterol compared to WT and *Prmt2^-/-^* baseline levels (baseline *Ldlr^-/-^:WT:* 885 mg/dL, baseline *Ldlr^-/-^:Prmt2^-/-^*: 862 mg/dL to 82-155 mg/dL in the regression groups). Also, the reduction in total cholesterol levels was independent of STZ treatment or the source of bone marrow (WT or *Prmt2^-/-^*; Sup. Fig. 1B; see Regression 3 weeks). As expected, the glucose levels were dramatically higher (~2.5-fold greater) in STZ-treated compared to control mice (Sup. Fig.1C; Regression 3 weeks D *Ldlr^-/-^:WT* and D *Ldlr^-/-^:Prmt2^-/-^*).

### Myeloid PRMT2 deficiency phenocopies the impairment in reducing plaque macrophage content in diabetes after lipid lowering

To determine the effect of myeloid PRMT2 status on atherosclerosis regression, plaque area and the contents (as % of plaque area) of macrophages (CD68+ cells) and collagen (Sirius red+ area under polarized light) were measured. Both groups of baseline mice had similar CD68+ (macrophage) plaque area and collagen content (Fig. 1 B-C; Sup. Fig. 2A-B). After hyperlipidemia was reduced, though plaque areas remained similar to baseline values (Fig. 1D), there were significant changes in plaque composition. This implies that as in other contexts, the effects of a given factor operate within a certain range of plasma cholesterol ^23^, with elevated levels being dominant over PRMT2-deficeincy. Reversal of hyperlipidemia, however, apparently allowed effects of PRMT2-deficiency to become penetrant, based on the changes in plaque composition. First, there was a significant reduction in plaque macrophage content in the NG *Ldlr^-/-^:WT* group. Although there was also a reduction in the NG *Ldlr^-/-^:Prmt2^-/-^* group, it was attenuated to a similar degree as we have observed in diabetic mice^17,23,24^, and as in the present study (compare the data in Fig. 1B between NG *Ldlr^-/-^:Prmt2^-/-^* and D *Ldlr^-/-^:WT*). In *Ldlr^-/-^:Prmt2^-/-^* mice, however, plaque macrophage content was similar in both normoglycemic and diabetic groups (Fig. 1B; Sup. Fig. 2A), suggesting that with myeloid deficiency of PRMT2, regression became independent of diabetes.

In contrast to the compositional changes, plaque areas (aortic root size) among the groups were not statistically significant (Fig. 1C). Consistent with our previous findings, the decreases in plaque macrophage content were mirrored by corresponding increases in collagen content (Fig. 1B; Sup. Fig. 2B), presumably because of less degradation of collagen by macrophages with lower level of activation (see below for data to support this presumption). The increases in collagen content were least in the NG *Ldlr^-/-^:Prmt2^-/-^*, D *Ldlr^-/-^:Prmt2^-/-^*, and D *Ldlr^-/-^:WT* groups, as expected from their more modest changes (vs. NG *Ldlr^-/-^:WT* mice) in plaque macrophage content.

As noted in the Introduction, hyperglycemia was associated with decreased expression of *Prmt2* in macrophages *in vitro*. To confirm that this was also true *in vivo*, in a separate set of mice, we measured *Prmt2* mRNA levels in plaque macrophages in normoglycemic and hyperglycemic mice. The levels of *Prmt2* mRNA in plaque macrophages (F4/80^+^ cells selected from aortic digestion by FACS) from diabetic mice were ~ 50% of those in normoglycemic.

Consistent with this are the data from myeloid cells in human plaques (originally reported in ^25^) in which the expression of *PRMT2* mRNA was statistically lower in samples taken from those with diabetes (Sup. Fig. 3). Together with the data in Figure 1, the results suggest that loss of PRMT2 expression either by genetic deficiency or by repression by hyperglycemia impairs the regression of plaque macrophage content after lipid lowering.

### PRMT2 deficiency increases macrophage retention, but does not change monocyte recruitment to, or macrophage proliferation in plaques

To investigate the basis for the difference in macrophage content among *Ldlr^-/-^:WT* and *Ldlr^-/-^:Prmt2^-/-^* regression groups, we examined key kinetic parameters that regulate plaque monocyte/macrophage abundance including recruitment, retention, proliferation, apoptosis and necrosis as we have done before^17,26–29^. The experimental design is shown Figure 2A. To assess monocyte recruitment during the regression period, mice were injected with fluorescent latex beads 3 days prior to harvest, and the number of beads in the aortic plaque was quantified and normalized to the efficiency of labeling circulating monocytes. The efficiency of monocyte labeling was not affected by either hyperglycemia or PRMT2-deficiency (not shown).

**Figure 2.**
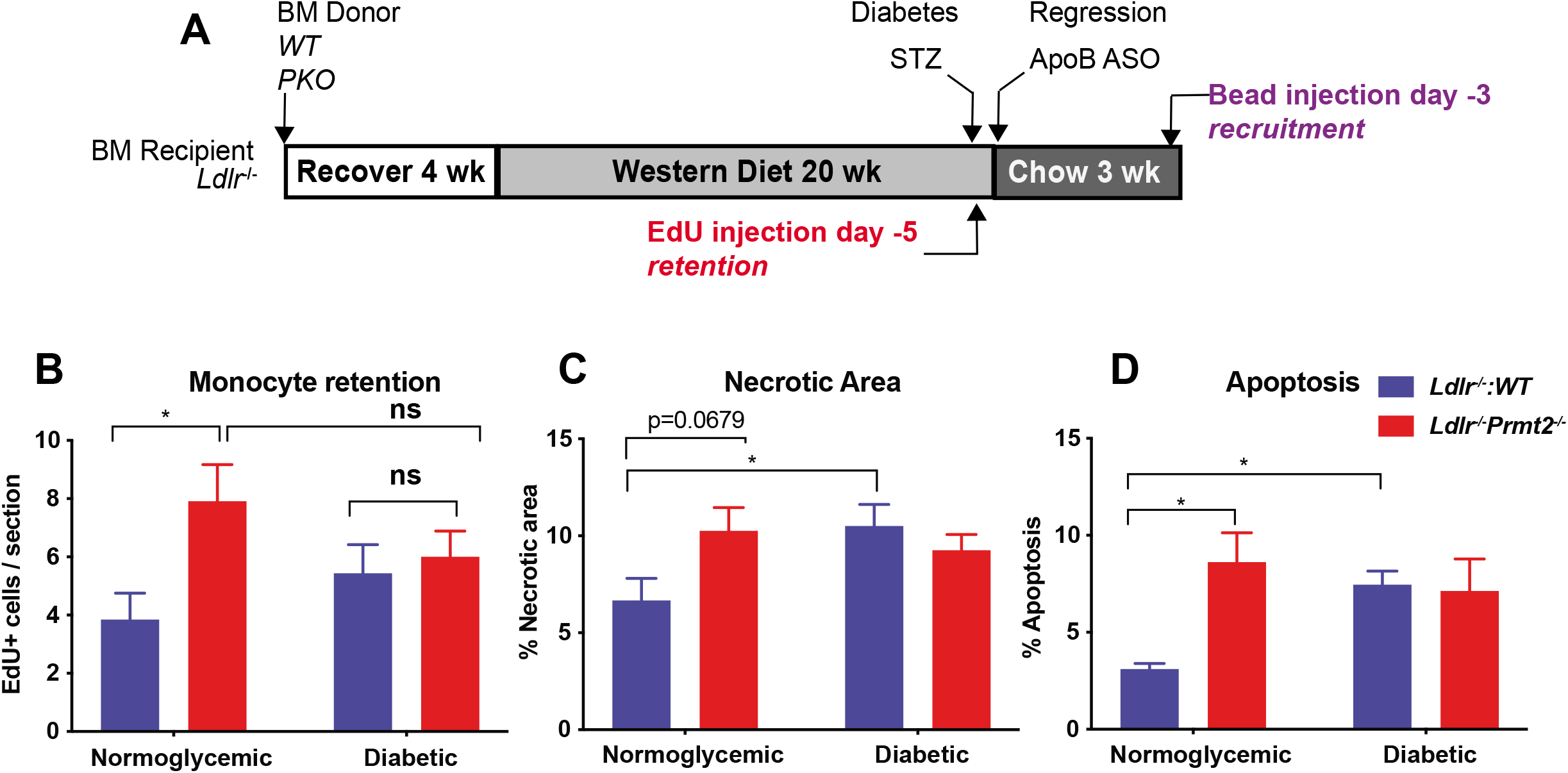
Macrophage dynamics show monocyte retention is the key kinetic change impairing regression in *Ldlr^-/-^:Prmt2^-/-^* mice in normoglycemia. A) Schematic of timeline for EdU and bead injections to assess retention and recruitment of monocytes, respectively, into aortic arches under regression conditions. B) Analysis of EdU+ cells/section of atherosclerotic plaques from the indicated groups showing significantly increased recruitment of EdU+ cells into *Ldlr^-/-^:Prmt2^-/-^* compared to *Ldlr^-/-^:WT* under normoglycemic conditions. C) Quantification of necrotic area and D) cleaved caspase 3 (to assess apoptosis) in plaque sections showing increased necrosis and apoptosis in *Ldlr^-/-^:Prmt2^-/-^* vs *Ldlr^-/-^:WT* under normoglycemic conditions. Data (means ± SEM) were analyzed using oneway ANOVA followed by Bonferroni’s multiple comparison test. *P* values are shown as \**P* < 0.05 and \*\*\*\**P* < 0.001.

Monocyte recruitment was not significantly different between the regression groups (NG *Ldlr^-/-^:WT*, NG *Ldlr^-/-^:Prmt2^-/-^*, D *Ldlr^-/-^:WT*, D *Ldlr^-/-^:Prmt2^-/-^*) (Sup. Fig. 4A). Diabetes did not significantly increase monocyte recruitment in WT mice (Sup. Fig. 4A). Thus, we turned to retention/emigration as a possible mechanism as the basis for the greater plaque macrophage content in the other 3 regression groups compared to the NG WT group.

To determine whether macrophage retention was relatively increased (reflecting a decreased capacity to emigrate) in the non-NG WT groups, mice were injected with EdU (5-ethynyl-2’-deoxyuridine) 5 days before the induction of regression, to mark cells that accumulate during plaque formation. We have previously reported^28^ that EdU preferentially labels circulating Ly6c^hi^ monocytes, thought to be the major contributors to the plaque macrophage pool^28^. The EdU labeling efficiency was similar between *Ldlr^-/-^:WT and Ldlr^-/-^:Prtm2^-/-^* baseline groups and averaged ~28 beads per section. Compared to the baseline groups, there was a decrease in EdU+ cells in all regression groups, suggesting that cells were actively leaving the plaque during regression. We found, however, that EdU+ macrophages were significantly more abundant in NG *Ldlr^-/-^:Prmt2^-/-^*, vs NG *Ldlr^-/-^:WT* mice (Fig. 2B). Furthermore, the EdU+ macrophage abundance in the NG *Ldlr^-/-^:Prmt2^-/-^* mice was not significantly different from either diabetes group (Fig. 2B).

To understand whether macrophage proliferation had an effect on plaque macrophage content, Ki67 (marker of cell proliferation) staining was performed (Sup. Fig. 4B). We observed that the number of Ki67 cells did not change among the regression groups, further supporting changes in macrophage retention/emigration as the basis for the increased CD68+ cell content in the NG *Ldlr^-/-^:Prmt2^-/-^* D *Ldlr^-/-^:Prmt2^-/-^*, and D *Ldlr^-/-^:WT* regression groups. This is in contrast to data in vascular injury models that PRMT2-deficiency promoted vascular smooth muscle cell proliferation ^30,31^, which likely reflects differential effects of PRMT2 by cell type or by perturbation. Furthermore, once myeloid cells are deficient in PRMT2, an additional effect of diabetes was not observed, implying that a major effect of hyperglycemia is mediated through its down-regulation of *Prmt2*.

### Effects of myeloid *Prmt2*-deficiency and diabetes on necrotic area and apoptosis

Besides macrophage content, another measure of disease severity in atherosclerosis is the size of the necrotic core, especially because of its association in humans with plaque rupture ^32,33^. Thus, we quantified the acellular areas (taken to reflect necrotic areas ^23^x) in aortic plaques. As shown in Figure 2C, the necrotic area was lowest in the NG *Ldlr^-/-^:WT* group. Notably the higher necrotic area in the NG *Ldlr^-/-^:Prmt2^-/-^* mice was similar to that in D *Ldlr^-/-^:Prmt2^-/-^* mice (as well as in the D *Ldlr^-/-^:WT* mice). As with the plaque macrophage content data (Fig. 1B) and retention data (Fig. 2B), diabetes did not exacerbate the effects of PRMT2-deficiency, again implying that another major effect of hyperglycemia is mediated through its down-regulation of PRMT2.

The acellular area is thought to result from apoptosis of macrophages. When we used caspase-3 staining to determine apoptosis, we found that the pattern paralleled the necrotic core area results (Fig. 2D), namely that the level of apoptosis was lowest in the NG *Ldlr^-/-^:WT* group, with similar (elevated) levels in the remaining 3 regression groups. Thus, the loss of PRMT2 expression by genetic deletion under normoglycemic conditions or upon reduced expression in diabetes promotes apoptosis resulting in a larger necrotic area. This effect in myeloid cells is in contrast to PRMT2-deficiency decreasing apoptosis in fibroblasts^12^, and, again, may reflect differential effects of cell type or perturbations.

### Loss of PRMT2 promotes expression of genes linked to cytokine signaling and inflammatory pathways in plaque CD68+ cells under normoglycemic conditions

To gain insight into why PRMT2-deficient myeloid cells fail to regress atherosclerosis compared to WT under non-diabetic conditions, we examined gene expression changes by performing bulk RNA sequencing (RNA seq) from NG *Ldlr^-/-^*:WT and NG *Ldlr^-/-^:Prmt2^-/-^* plaque CD68+ cells collected by laser-capture microdissection^20^. We found in non-diabetic conditions CD68+ *Ldlr^-/-^:Prmt2^-/-^* cells induced 204 and repressed 120 genes compared to *Ldlr^-/-^*:WT (LogFC < 2.0, *P* < 0.01) (Fig. 3A). Metascape^21^ pathway analysis indicates that the up-regulated genes in *Ldlr^-/-^:Prmt2^-/-^* CD68+ cells are involved with cytokine signaling and inflammatory/innate immune responses (Fig. 3B). Moreover, analysis of the interactions between transcription factors (TFs) and the gene targets using the TRRUST (transcriptional regulatory relationships unraveled by sentence-based text-mining) algorithm^34^ identified NF-kappa B, followed by IRF1 and STAT1as the top TFs controlling gene expression in *Ldlr^-/-^:Prmt2^-/-^* vs *Ldlr^-/-^:WT* CD68+ cells under normoglycemic conditions (Fig. 3C). The top GO class associated with the genes repressed in *Ldlr^-/-^:Prmt2^-/-^* compared to *Ldlr^-/-^:WT* CD68+ cells under normoglycemia is the regulation of pseudopodium assembly (Sup. Fig. 5A). Pseudopodia are used by macrophages for phagocytosis^35^. The top transcription factor associated with the genes reduced upon *Prmt2* deletion is EGR1 (Sup. Fig. 5B). EGR1 has been shown to occupy enhancers associated with inflammatory response genes to increase their expression^36^. Thus, RNA sequencing reveals a heightened inflammatory state in PRMT2-deficient plaque macrophages under non-diabetic conditions.

**Figure 3.**
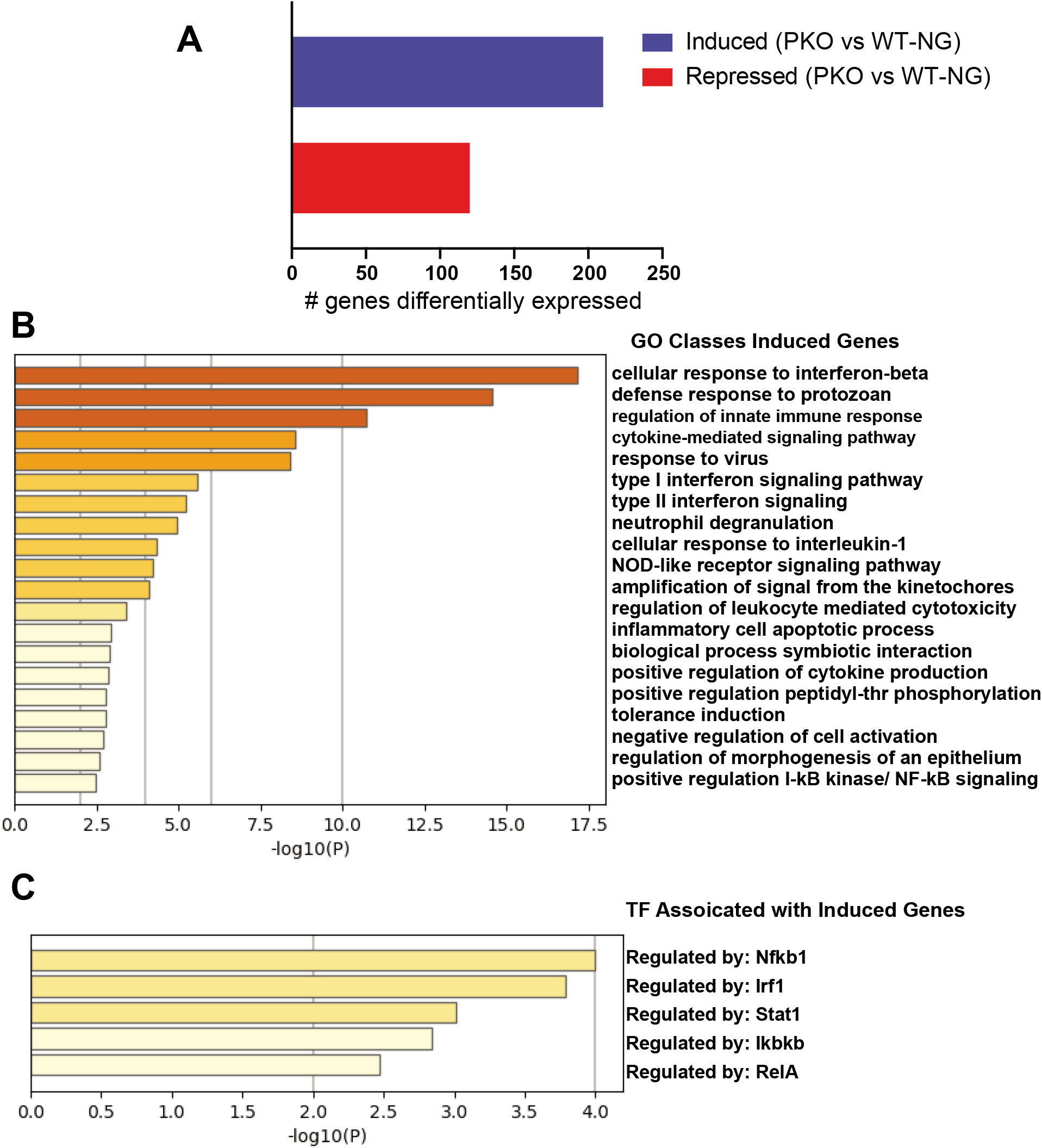
GO classes and transcription factors associated with genes up-regulated in *Ldlr^-/-^:Prmt2^-/-^* plaque CD68+ cells under normoglycemic conditions. A) PRMT2-deficiency in plaque CD68+ cells resulted in the up-regulation of 204 and down-regulation of 120 genes (LogFC < 2.0, *P* < 0.01). B) Metascape analysis of pathways for genes up-regulated in *Ldlr^-/-^:Prmt2^-/-^* reveals genes involved cytokine signaling and innate immune/inflammatory responses. C) Transcription factors associated with the genes up-regulated in *Ldlr^-/-^:Prmt2^-/-^* in normoglycemia include the prototypical proinflammatory transcriptional regulatory factors NF-kappa B, IRF1 and STAT1.

That the RNA sequencing results showed that PRMT2-deficient plaque macrophages had a more inflammatory transcriptional profile is consistent with its reported role in regulating the inflammatory responses^12,13^. Specifically, in fibroblasts PRMT2 has been shown to be an inhibitor of NF-kappa B and its deficiency dampens lung macrophage responsiveness to LPS^12,13^. To investigate whether PRMT2 plays similar roles in our system, BMDMs were isolated from either *Prmt2*-sufficient or deficient mice and maintained during their 7 day differentiation period under metabolic conditions to simulate normoglycemia (5 mM D-glucose) or hyperglycemia (25 mM D-glucose). As a control for the osmolality difference, the medium that was 5mM D-glucose was supplemented with 20 mM L-glucose. Cells were then treated with either IL-4 to promote M2 polarization, or LPS to induce the M1 state. We then measured the expression of standard markers of macrophages responding to inflammation resolving IL-4 (*Arg1* and *Fizz1*) and inflammation promoting LPS (*Il6* and *Nos2*) as a function of normal glucose and high glucose, and in either the absence or present of PRMT2.

We had previously reported that responsiveness to IL-4, which promotes inflammation resolution, is required for plaque regression^28^, and that hyperglycemia attenuates macrophage responsiveness to IL-4^17^. Whereas the induction *Arg1* and *Fizz1* by IL-4 is blunted, as expected, by hyperglycemia, the effect of PRMT2-deficiency in BMDMs is similar to that of hyperglycemia in that the levels of *Arg1* and *Fizz1* are lower compared to the corresponding values for the sufficient cells (Fig. 4 A-B). Thus, the loss of PRMT2 in normal glucose mimics the high glucose state.

**Figure 4.**
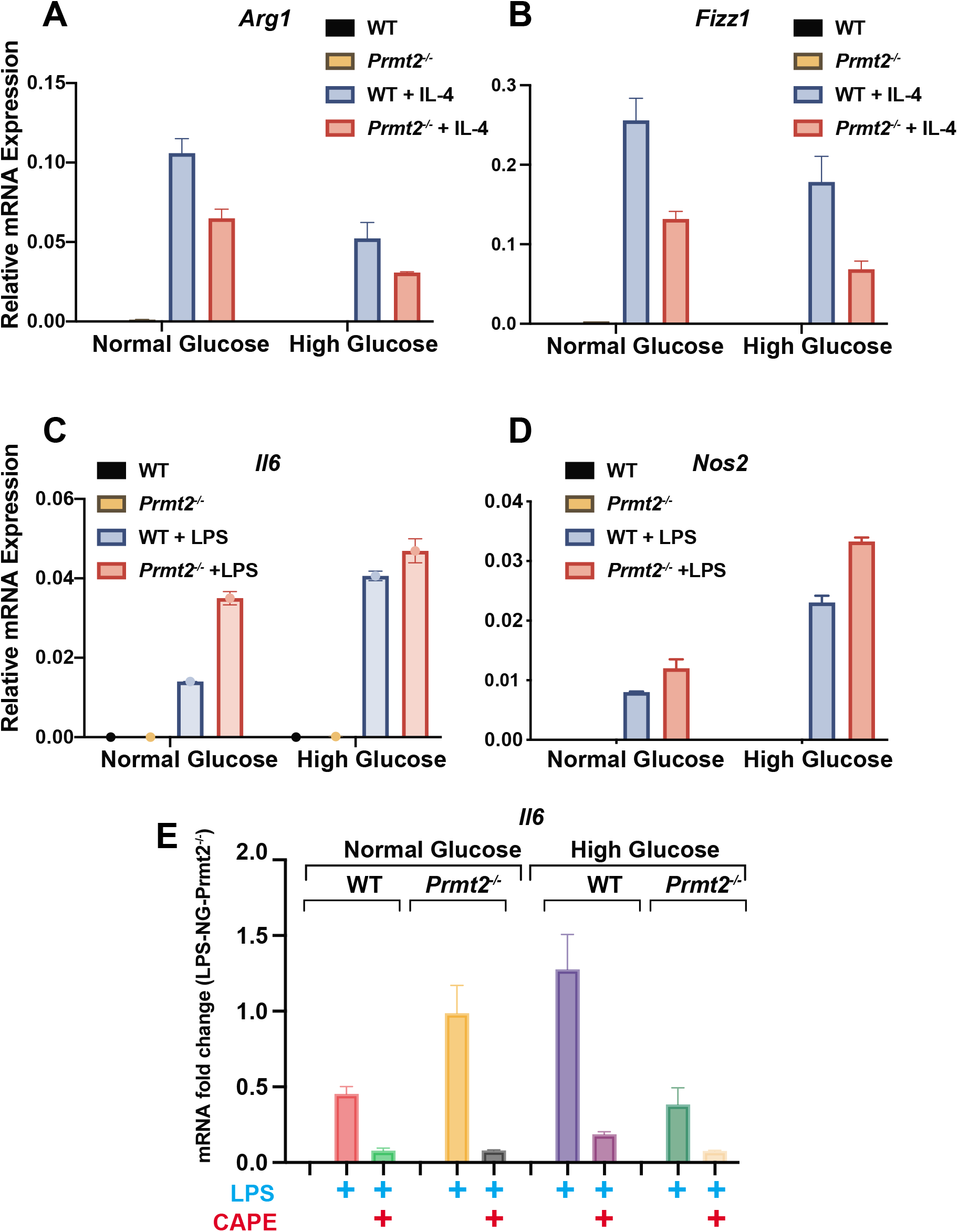
PRMT2-deficeny blunts macrophage response by IL-4 and heightens macrophage response to LPS *in vitro*. BMDMs from WT and *Prmt2^-/-^* mice were differentiated into M0 (unactivated) macrophages in normal d+l-glucose (100 mg/dL d-glucose + 350 mg/dL l-glucose) or high d-glucose (450 mg/dL) for 7 days, and then treated with or without IL-4 (to induce M2 polarization) or with or without LPS (to induce M1 polarization). RNA was prepared and qPCR was performed on M2 marker genes A) arginase 1 (*Arg1*) and B) *Fizz1*, and M1 markers C) *Il6* and D) *Nos2*, and normalized to cyclophilin A. E) LPS induction of *Il6* is NF-kappa B dependent. BMDMs were differentiated in normal or high glucose as above. Cells were pretreated for 2 hours with NF-kappa B inhibitor CAPE (5 μM) and then treated with LPS for 6 hours. RNA was isolated and *Il6* expression was determined by qPCR relative to cyclophilin A1 and shown as fold change with normal glucose *Prmt2^-/-^* mRNA set to 1.0. The data presented are means with error bars representing the range of the means from an experiment that was repeated twice.

As noted above, PRMT2 is a known inhibitor of NF-kappa B with a mechanism involving blocking nuclear export of IkB-alpha, resulting in decreased NF-kappa B binding to its transcriptional targets^12^.Thus, we hypothesize that the increases in the expression of targets of NF-kappa B would result from the relief of this inhibition. To test this we stimulated WT and *Prmt2^-/-^* BMDMs with LPS and measured the expression of the NF-kappa B target genes *Il6* and *Nos2*. We found that high glucose compared to normal glucose increased *Il6* and *Nos2* expression and this was further amplified in *Prmt2^-/-^* compared to WT BMDMs (Fig. 4 C-D). To demonstrate a direct role of NF-kappa B in the heightened inflammatory state of PRMT2-deficient BMDMs, we also examined whether the induction of *Il6* by LPS was reduced upon inhibition of NF-kappa B by caffeic acid phenethyl ester (CAPE), an inhibitor that blocks NF-kappa B binding to DNA^37^, or TPCA1^38,39^, a potent and selective inhibitor the IκB kinase 2 (IKK-2) required for NF-kappa B activation^40^. When either NF-kappa B inhibitor was used the increase in *Il6* under normal and high glucose in WT and *Prmt2^-/-^* BMDMs was attenuated (Fig. 4E: Sup. Fig. 6). Taken together, these data are consistent with the RNA-seq analysis that suggested that PRMT2-deficiency heightens the inflammatory state of plaque macrophages, and suggests that the acquisition of inflammatory phenotype by the loss of PRMT2 is through activation of NF-kappa B. Furthermore, PRMT2-deficiency not only increases responsiveness to an inflammatory stimulus, it also blunts the response of macrophages to the potent inflammation resolver, IL-4.

## CONCLUDING REMARKS

Diabetes impairs the benefits of cholesterol-lowering clinically^1^, and we have previously shown consistent results in a mouse model of atherosclerosis regression^17,23,24,41^. When probing the molecular basis for these adverse effects of diabetes, we found the elevated glucose represses the expression of PRMT2^6^, leading to the present studies to test the hypothesis that part of the effects of diabetes was a consequence of this repression. Indeed, the data presented support this hypothesis, particularly by the impairment in the desirable remodeling of the plaque composition after cholesterol loading in mice with myeloid deficiency of PRMT2 that were normoglycemic. The ability of PRMT2-deficiency to phenocopy effects of diabetes in the regression, but not progression, setting implies that it would be more of an important factor in people undergoing risk factor reduction than during the unabated progression of their disease. The most plausible molecular mechanism for the impaired resolution of inflammation, required for maximal responses to cholesterol lowering in mice ^28^ and people (CANTOS)^42^, is the inhibition of NF-kappa B by PRMT2. The increasing availability of curated data sets from human atherosclerotic plaques from people with and without diabetes will be a valuable resource with which to extend our mouse model findings to the clinical scenario. Indeed, lower PRMT2 expression was observed in myeloid cells of plaques from humans with diabetes compared to those without, supporting the relevance to our findings in mouse models to human atherosclerosis.

## Abbreviations

PRMT2: Protein Arginine Methyltransferase 2
Ldlr: low density lipoprotein receptor
CHD: coronary heart disease
LXR: liver X receptor
ASO: anti-sense oligonucleotide
APOB: apolipoprotein B

---

**Supplementary Figure 1. Plasma cholesterol concentrations, blood glucose levels and weights from *Ldlr^-/-^:WT* and *Ldlr^-/-^:Prmt2^-/-^* cohorts**

A) Plasma total cholesterol concentrations, B) glucose levels and C) body weight were measured at the 19-week time point of western diet feeding for the baseline groups, and at the end of the experiment for the indicated regression groups under conditions of normoglycemia and diabetes.

**Supplementary Figure 2. Images of CD68 and collagen content in plaques from *Ldlr^-/-^:WT* and *Ldlr^-/-^:Prmt2^-/-^* with and without diabetes**

Representative micrographs of A) CD68+ staining and B) collagen content as determined by picosirius red staining and imaged using brightfield and polarized light microscopy.

**Supplementary Figure 3. Expression of PRMT2 in myeloid cells from human atherosclerotic plaques.**

Single-cell RNA sequencing of myeloid cells from human atherosclerotic plaques from nondiabetic and diabetic (type 2) patients was analyzed for expression of *PRMT2* (N=6). Data were acquired from sequence information in Fernandez *et al* ^25^.

**Supplementary Figure 4. Monocyte recruitment and macrophage proliferation in plaques are not significantly different between *Ldlr^-/-^:WT* and *Ldlr^-/-^:Prmt2^-/-^* in normoglycemic and diabetic conditions**

A) Analysis of bead^+^ cells/section of atherosclerotic plaques are shown for each group. B) Quantification in plaque sections of Ki67 to assess proliferation. No significant differences were observed in either monocyte recruitment or proliferation between recipient groups.

**Supplementary Figure 5. GO classes and transcription factor associated with genes down-regulated in PRMT2-deficient plaque CD68+ cells in normoglycemic conditions**

A) Metascape analysis of the120 genes down-regulated genes in *Ldlr^-/-^:Prmt2^-/-^* plaque CD68+ cells compared to *Ldlr^-/-^:WT* reveals pathways involved in pseudopodium assembly among others. B) Transcription factor EGR1 is associated with the genes down-regulated in *Ldlr^-/-^:Prmt2^-/-^* in normoglycemia.

**Supplementary Figure 6. LPS induction of *Il6* expression is blocked by the IκB kinase 2 (IKK-2) inhibitor TPCA1**

WT and *Prmt2^-/-^* BMDMs were differentiated in normal or high glucose. Cells were pretreated for 2 hours with NF-kappa B inhibitor TPCA1 (5 μM) and then treated with LPS for 6 hours. RNA was isolated and *Il6* expression was determined by qPCR relative to cyclophilin A1 and shown as fold change with normal glucose *Prmt2^-/-^* mRNA set to 1.0. The data presented are means ± standard errors of the means of three independent experiments.

## ACKNOWLEDGEMENTS

We thank Drs. Elizabeth Nabel and Yaan Herault for generously providing the PRMT2^-/-^ mice. This work was supported by DOD grant W81XWH-16-1-0373 (EAF, MJG).

## REFERENCES

1 Fox, C. S. et al. Increasing cardiovascular disease burden due to diabetes mellitus: the Framingham Heart Study. Circulation 115, 1544–1550, doi:10.1161/CIRCULATIONAHA.106.658948 (2007).

2 Moore, K. J., Sheedy, F. J. & Fisher, E. A. Macrophages in atherosclerosis: a dynamic balance. Nat Rev Immunol 13, 709–721, doi:10.1038/nri3520 (2013).

3 Nagareddy, P. R. et al. Hyperglycemia promotes myelopoiesis and impairs the resolution of atherosclerosis. Cell Metab 17, 695–708, doi:10.1016/j.cmet.2013.04.001 (2013).

4 Rahman, K. & Fisher, E. A. Insights From Pre-Clinical and Clinical Studies on the Role of Innate Inflammation in Atherosclerosis Regression. Front Cardiovasc Med 5, 32,doi:10.3389/fcvm.2018.00032 (2018).

5 Williams, K. J., Feig, J. E. & Fisher, E. A. Rapid regression of atherosclerosis: insights from the clinical and experimental literature. Nat Clin Pract Cardiovasc Med 5, 91–102, doi:10.1038/ncpcardio1086 (2008).

6 Hussein, M. A. et al. LXR-Mediated ABCA1 Expression and Function Are Modulated by High Glucose and PRMT2. PloS one 10, e0135218,doi:10.1371/journal.pone.0135218 (2015).

7 Dong, F. et al. PRMT2 links histone H3R8 asymmetric dimethylation to oncogenic activation and tumorigenesis of glioblastoma. Nat Commun 9, 4552,doi:10.1038/s41467-018-06968-7 (2018).

8 Blythe, S. A., Cha, S. W., Tadjuidje, E., Heasman, J. & Klein, P. S. beta-Catenin primes organizer gene expression by recruiting a histone H3 arginine 8 methyltransferase, Prmt2. Dev Cell 19, 220–231, doi:10.1016/j.devcel.2010.07.007 (2010).

9 Wu, Q., Schapira, M., Arrowsmith, C. H. & Barsyte-Lovejoy, D. Protein arginine methylation: from enigmatic functions to therapeutic targeting. Nat Rev Drug Discov 20, 509–530, doi:10.1038/s41573-021-00159-8 (2021).

10 Blanc, R. S. & Richard, S. Arginine Methylation: The Coming of Age. Mol Cell 65, 8–24, doi:10.1016/j.molcel.2016.11.003 (2017).

11 Cura, V. & Cavarelli, J. Structure, Activity and Function of the PRMT2 Protein Arginine Methyltransferase. Life (Basel) 11, doi:10.3390/life11111263 (2021).

12 Ganesh, L. et al. Protein methyltransferase 2 inhibits NF-kappaB function and promotes apoptosis. Mol Cell Biol 26, 3864–3874, doi:10.1128/MCB.26.10.3864-3874.2006 (2006).

13 Dalloneau, E., Pereira, P. L., Brault, V., Nabel, E. G. & Herault, Y. Prmt2 regulates the lipopolysaccharide-induced responses in lungs and macrophages. J Immunol 187, 4826–4834, doi:10.4049/jimmunol.1101087 (2011).

14 Peet, D. J., Janowski, B. A. & Mangelsdorf, D. J. The LXRs: a new class of oxysterol receptors. Curr Opin Genet Dev 8, 571–575, doi:10.1016/s0959-437x(98)80013-0 (1998).

15 Wu, C. et al. Modulation of Macrophage Gene Expression via Liver X Receptor alpha Serine 198 Phosphorylation. Mol Cell Biol 35, 2024–2034, doi:10.1128/MCB.00985-14 (2015).

16 Feig, J. E. et al. LXR promotes the maximal egress of monocyte-derived cells from mouse aortic plaques during atherosclerosis regression. The Journal of clinical investigation 120, 4415–4424, doi:10.1172/JCI38911 (2010).

17 Parathath, S. et al. Diabetes adversely affects macrophages during atherosclerotic plaque regression in mice. Diabetes 60, 1759–1769, doi:10.2337/db10-0778 (2011).

18 Choudhury, R. P. et al. High-density lipoproteins retard the progression of atherosclerosis and favorably remodel lesions without suppressing indices of inflammation or oxidation. Arterioscler Thromb Vasc Biol 24, 1904–1909, doi:10.1161/01.ATV.0000142808.34602.25 (2004).

19 Junqueira, L. C., Bignolas, G. & Brentani, R. R. Picrosirius staining plus polarization microscopy, a specific method for collagen detection in tissue sections. Histochem J 11, 447–455, doi:10.1007/BF01002772 (1979).

20 Feig, J. E. & Fisher, E. A. Laser capture microdissection for analysis of macrophage gene expression from atherosclerotic lesions. Methods Mol Biol 1027, 123–135, doi:10.1007/978-1-60327-369-5_5 (2013).

21 Zhou, Y. et al. Metascape provides a biologist-oriented resource for the analysis of systems-level datasets. Nat Commun 10, 1523, doi:10.1038/s41467-019-09234-6 (2019).

22 Sharma, M. et al. Regulatory T Cells License Macrophage Pro-Resolving Functions During Atherosclerosis Regression. Circulation research 127, 335–353, doi:10.1161/CIRCRESAHA.119.316461 (2020).

23 Distel, E. et al. miR33 inhibition overcomes deleterious effects of diabetes mellitus on atherosclerosis plaque regression in mice. Circulation research 115, 759–769, doi:10.1161/CIRCRESAHA.115.304164 (2014).

24 Willecke, F. et al. Effects of High Fat Feeding and Diabetes on Regression of Atherosclerosis Induced by Low-Density Lipoprotein Receptor Gene Therapy in LDL Receptor-Deficient Mice. PloS one 10, e0128996, doi:10.1371/journal.pone.0128996 (2015).

25 Fernandez, D. M. et al. Single-cell immune landscape of human atherosclerotic plaques. Nat Med 25, 1576–1588, doi:10.1038/s41591-019-0590-4 (2019).

26 Feig, J. E., Hewing, B., Smith, J. D., Hazen, S. L. & Fisher, E. A. High-density lipoprotein and atherosclerosis regression: evidence from preclinical and clinical studies. Circulation research 114, 205–213, doi:10.1161/CIRCRESAHA.114.300760 (2014).

27 Feig, J. E. et al. Regression of atherosclerosis is characterized by broad changes in the plaque macrophage transcriptome. PloS one 7, e39790, doi:10.1371/journal.pone.0039790 (2012).

28 Rahman, K. et al. Inflammatory Ly6Chi monocytes and their conversion to M2 macrophages drive atherosclerosis regression. The Journal of clinical investigation 127, 2904–2915, doi:10.1172/JCI75005 (2017).

29 Weinstock, A. & Fisher, E. A. Methods to Study Monocyte and Macrophage Trafficking in Atherosclerosis Progression and Resolution. Methods Mol Biol 1951, 153–165, doi:10.1007/978-1-4939-9130-3_12 (2019).

30 Zeng, S. Y. et al. Protein Arginine Methyltransferase 2 Inhibits Angiotensin II-Induced Proliferation and Inflammation in Vascular Smooth Muscle Cells. Biomed Res Int 2018, 1547452,doi:10.1155/2018/1547452 (2018).

31 Yoshimoto, T. et al. The arginine methyltransferase PRMT2 binds RB and regulates E2F function. Exp Cell Res 312, 2040–2053, doi:10.1016/j.yexcr.2006.03.001 (2006).

32 Bentzon, J. F., Otsuka, F., Virmani, R. & Falk, E. Mechanisms of plaque formation and rupture. Circulation research 114, 1852–1866, doi:10.1161/CIRCRESAHA.114.302721 (2014).

33 Zhu, S. N., Chen, M., Jongstra-Bilen, J. & Cybulsky, M. I. GM-CSF regulates intimal cell proliferation in nascent atherosclerotic lesions. J Exp Med 206, 2141–2149, doi:10.1084/jem.20090866 (2009).

34 Han, H. et al. TRRUST v2: an expanded reference database of human and mouse transcriptional regulatory interactions. Nucleic Acids Res 46, D380–D386, doi:10.1093/nar/gkx1013 (2018).

35 Aderem, A. & Underhill, D. M. Mechanisms of phagocytosis in macrophages. Annu Rev Immunol 17, 593–623, doi:10.1146/annurev.immunol.17.1.593 (1999).

36 Trizzino, M. et al. EGR1 is a gatekeeper of inflammatory enhancers in human macrophages. Sci Adv 7, doi:10.1126/sciadv.aaz8836 (2021).

37 Natarajan, K., Singh, S., Burke, T. R., Jr., Grunberger, D. & Aggarwal, B. B. Caffeic acid phenethyl ester is a potent and specific inhibitor of activation of nuclear transcription factor NF-kappa B. Proceedings of the National Academy of Sciences of the United States of America 93, 9090–9095, doi:10.1073/pnas.93.17.9090 (1996).

38 Podolin, P. L. et al. Attenuation of murine collagen-induced arthritis by a novel, potent, selective small molecule inhibitor of IkappaB Kinase 2, TPCA-1 (2-[(aminocarbonyl)amino]-5-(4-fluorophenyl)-3-thiophenecarboxamide), occurs via reduction of proinflammatory cytokines and antigen-induced T cell Proliferation. J Pharmacol Exp Ther 312, 373–381, doi:10.1124/jpet.104.074484 (2005).

39 Baxter, A. et al. Hit-to-lead studies: the discovery of potent, orally active, thiophenecarboxamide IKK-2 inhibitors. Bioorg Med Chem Lett 14, 2817–2822, doi:10.1016/j.bmcl.2004.03.058 (2004).

40 Gamble, C. et al. Inhibitory kappa B Kinases as targets for pharmacological regulation. Br J Pharmacol 165, 802–819, doi:10.1111/j.1476-5381.2011.01608.x (2012).

41 Barrett, T. J., Murphy, A. J., Goldberg, I. J. & Fisher, E. A. Diabetes-mediated myelopoiesis and the relationship to cardiovascular risk. Ann N Y Acad Sci 1402, 31–42, doi:10.1111/nyas.13462 (2017).

42 Ridker, P. M. et al. Antiinflammatory Therapy with Canakinumab for Atherosclerotic Disease. N Engl J Med 377, 1119–1131, doi:10.1056/NEJMoa1707914 (2017).

