## Supplementary material for "Loss of PRMT2 in myeloid cells in normoglycemic mice phenocopies impaired regression of atherosclerosis in diabetic mice": Supplmental Figures 1-6

### Supplementary Figure 1

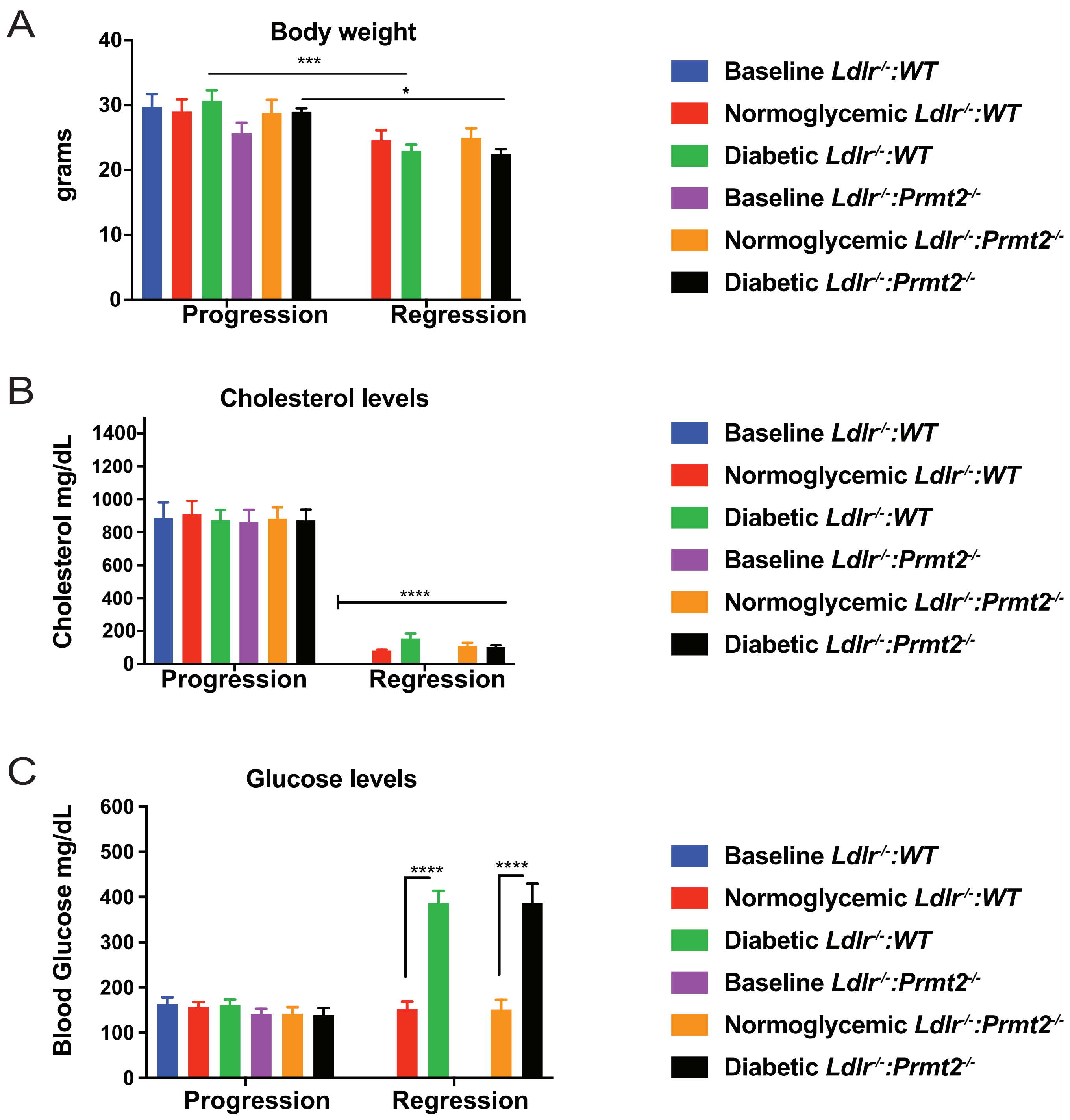

### Supplementary Figure 2

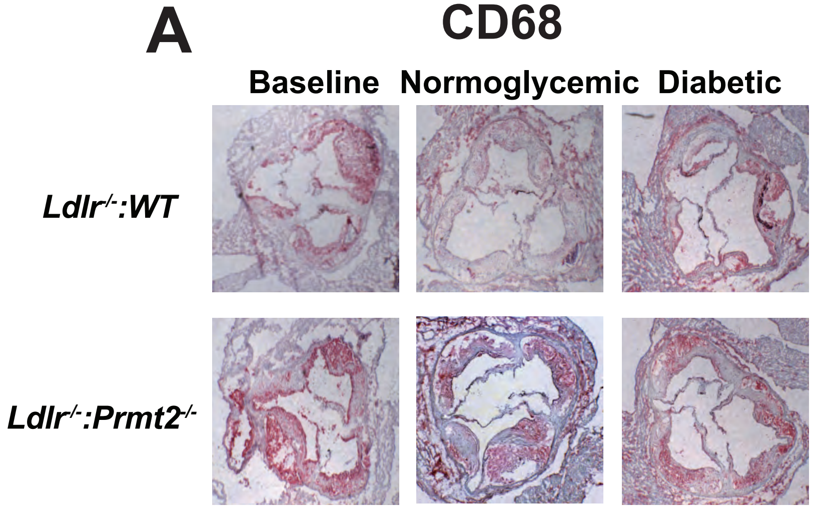


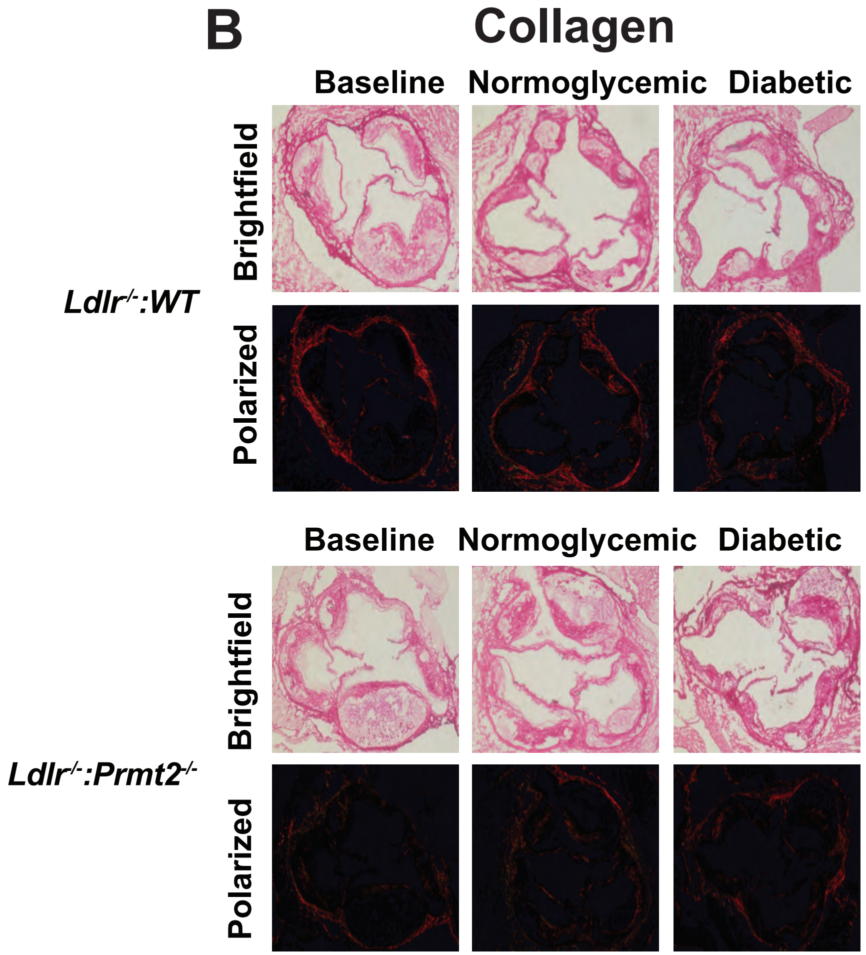

### Supplementary Figure 3

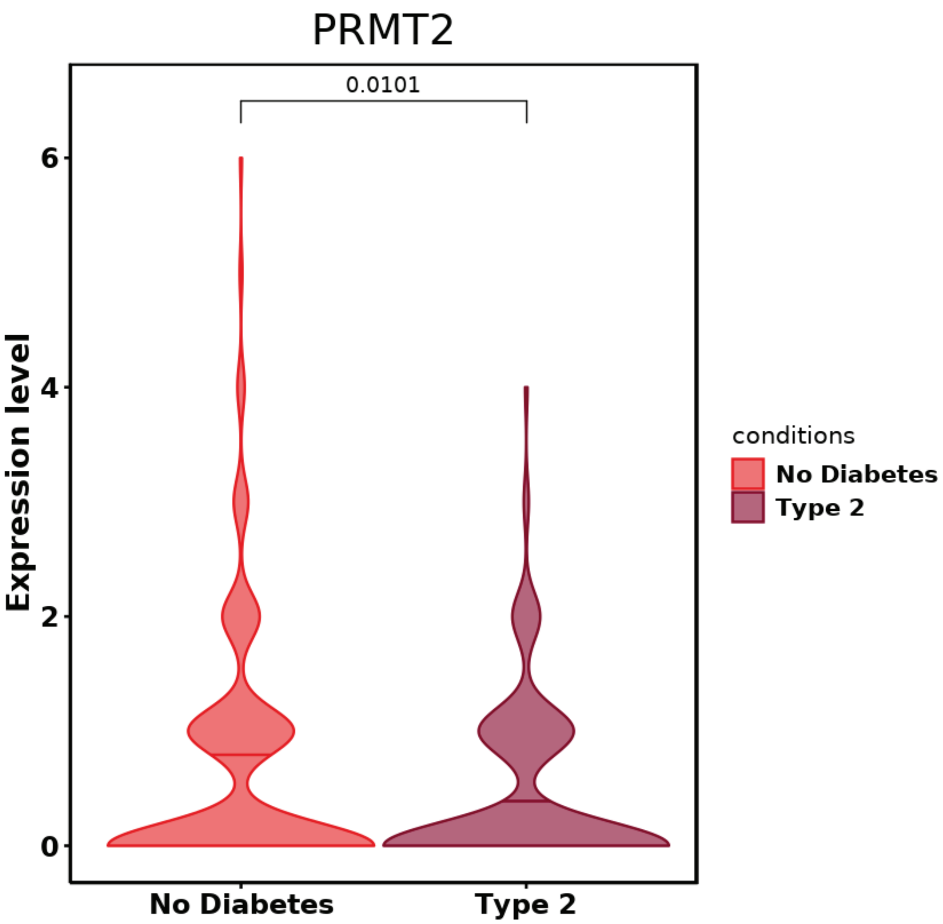

### Supplementary Figure 4

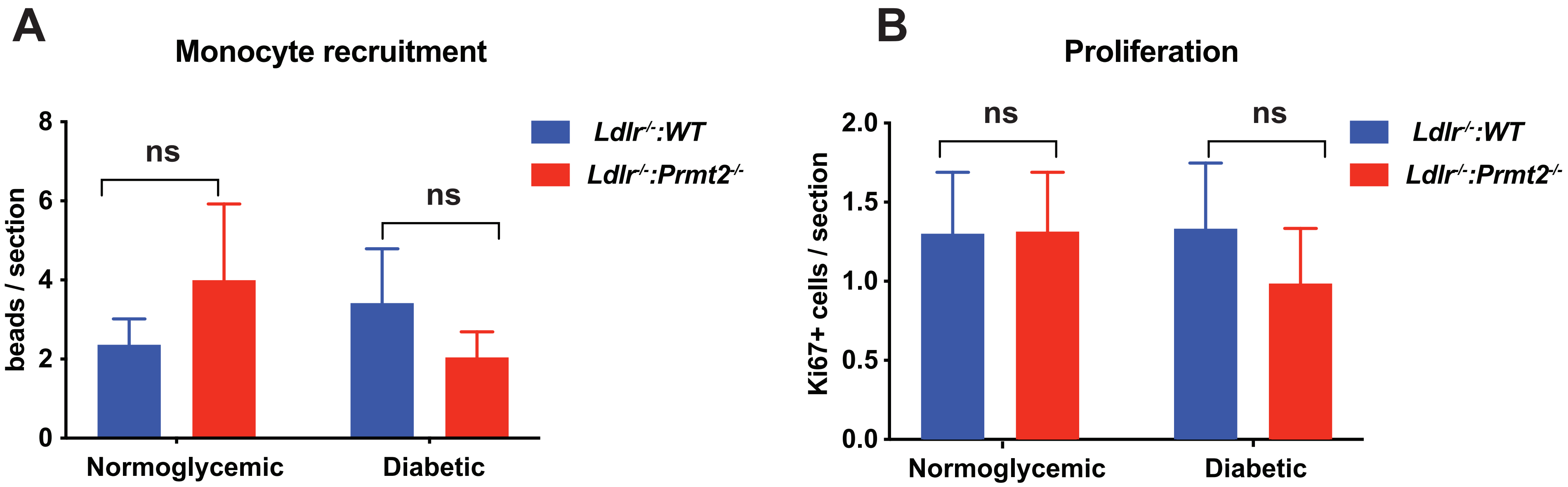

### Supplementary Figure 5

**A**

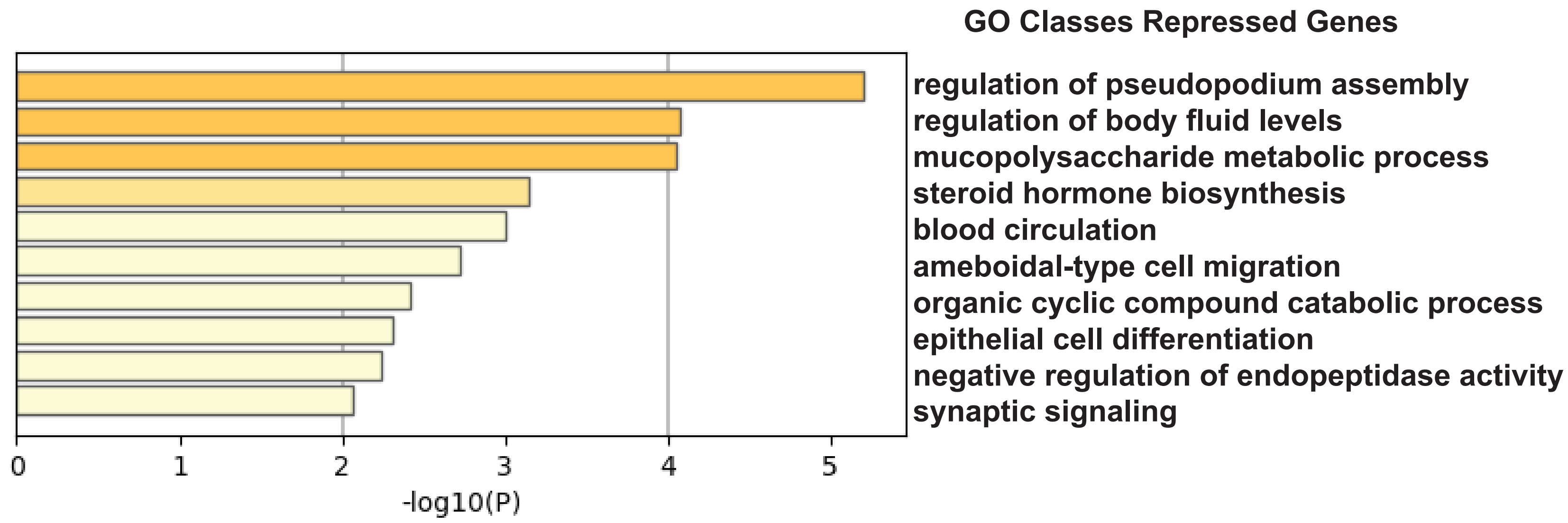


**B**

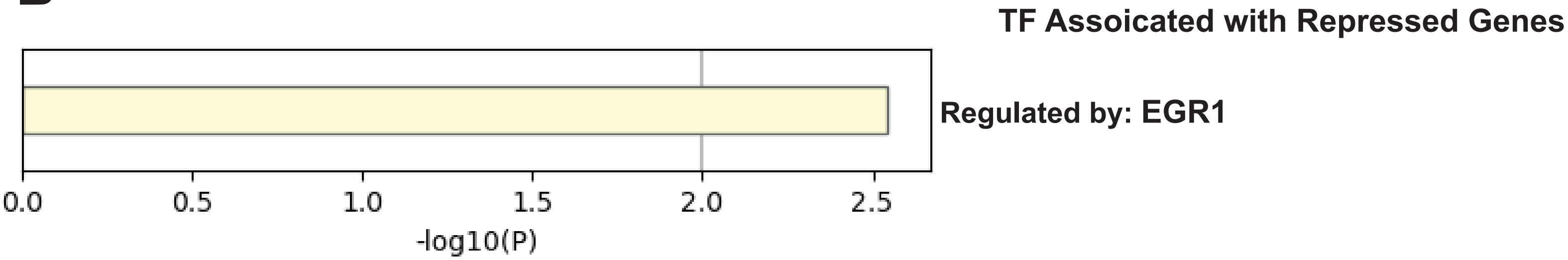

### Supplementary Figure 6

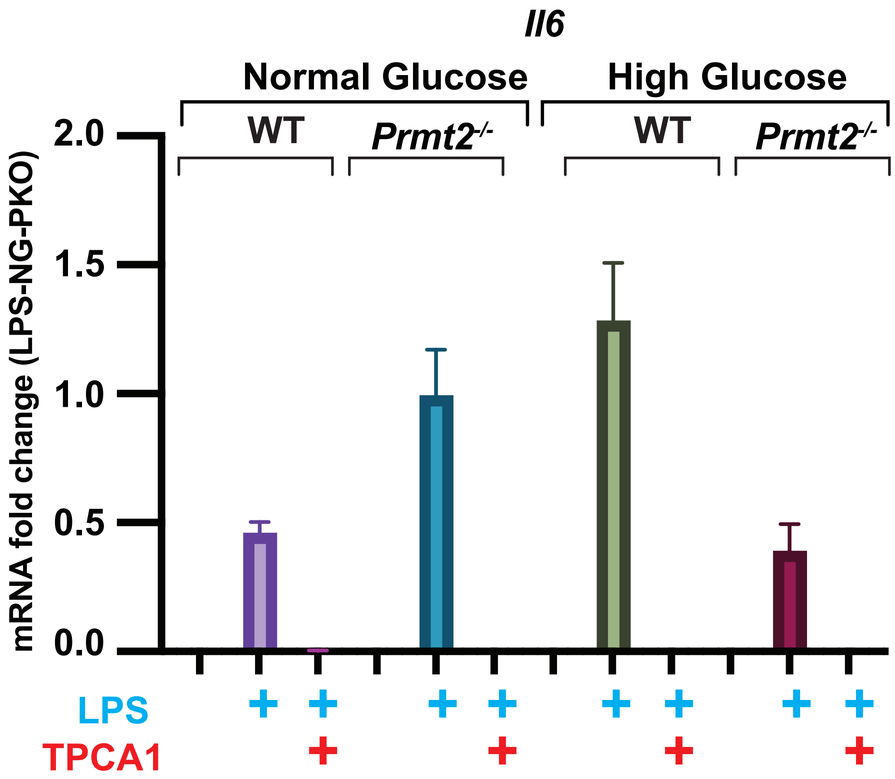
